# Mesodermal-specific *MECP2* expression in *Drosophila* induces visceral and skeletal muscle defects rescued by butyrate supplementation

**DOI:** 10.64898/2025.12.16.694631

**Authors:** Gaia Consonni, Mustafa Kaan Kozluca, Maria Cristina Gagliani, Alessandro Mineo, Katia Cortese, Irene Miguel-Aliaga, Aglaia Vignoli, Elisa Borghi, Antonio Galeone, Thomas Vaccari

## Abstract

**Background:** Patients affected by Rett syndrome (RTT) and MECP2 duplication syndrome (MDS) experience disabling muscle weakness and gastrointestinal dysmotility of unclear origin. Whether these defects arise cell-autonomously, rather than secondarily to neural dysfunction, and which developmental windows are most vulnerable to MeCP2 disfunction remains unresolved. MeCP2 is a dosage-sensitive transcriptional regulator, whose functions are tightly linked to chromatin states. Because short-chain fatty acids (SCFAs) are known to inhibit histone deacetylases (HDACs), a tractable *in vivo* model is needed to test the effect of HDAC modulation on muscle defects.

**Methods:** We misexpressed human *MECP2* in the *Drosophila melanogaster* mesoderm that gives rise to skeletal and visceral muscles. We analyzed quantitatively their morphology and function. To assess the effects of SCFA supplementation, we also supplemented diets with sodium butyrate (NaB), Lalbaay®, a NaB-containing supplement, acetate (AcOH), and valproate (VPA).

**Findings:** *MECP2* misexpression caused pre-eclosion lethality, thinning of larval skeletal fibers with nuclear mispositioning and altered mitochondria. Functionally, it reduced locomotion, decreased food transit and gut peristalsis. Phenotypes were strongest when expression began during development. NaB and VPA supplementation rescue most of these phenotypes, consistent with their histone-deacetylase (HDAC) activity. Defects were not observed upon comparable misexpression of an RTT-associated MeCP2 loss-of-function variant, indicating that they might be relevant to pathogenesis of MECP2-related disorders.

**Interpretation:** Our genetic *in vivo* analysis models peripheral effects of MeCP2 dysregulation and their amelioration, supporting the possibility of HDAC-targeted strategies for MECP2-related muscle and gastrointestinal dysfunction.

## Introduction

Methyl-CpG-binding protein 2 (MeCP2) is an epigenetic regulator of chromatin, which interacts with methylated DNA and remodeling complexes (Amir et al., 1999; Collins & Neul, 2022; Samaco & Neul, 2011). Beyond its canonical affinity for methylated CpG islands, MeCP2 also binds unmethylated DNA structures and can modulate chromatin independently of DNA methylation (Galvão & Thomas, 2005; Hansen et al., 2010; Yakabe et al., 2008). The protein contains multiple functional domains, including the methyl-CpG-binding domain (MBD) and the transcriptional repression domain (TRD), which regulate chromatin accessibility and transcription of MeCP2 target genes (Samaco & Neul, 2011).

RTT and MDS are distinct X-linked developmental disorders caused by loss-of-function and gain-of-function mutations in *MECP2*, respectively (Amir et al., 1999; Van Esch et al., 2005; Yamamoto et al., 2014). Despite opposite dosage perturbations, the two disorders share overlapping clinical features, including seizures, motor impairments, and gastrointestinal (GI) dysfunction (Ihekweazu & Motil, 2025; Peron et al., 2022; Allison et al., 2024).

The GI manifestations in RTT are pervasive and debilitating. In a nationwide caregiver survey of 118 females with RTT, 81% reported GI symptoms “sometimes, often, or almost always,” with constipation (81%), gas and bloating (70%), and difficulties with eating, chewing, or swallowing (73%) most common (Ihekweazu & Motil, 2025). Nearly half noted frequent irritability or discomfort linked to GI issues, and over 60% required laxatives. These problems contribute to malnutrition, growth failure, and reduced quality of life, highlighting the need for targeted interventions (Ihekweazu & Motil, 2025). Musculoskeletal and nutritional-related complications are likewise prominent. A recent systematic review and meta-analysis reported scoliosis in ∼65% of patients (rising to 73–80% in adolescents), foot deformities in ∼53%, and hip displacement in ∼30%, while low bone density and fracture risk are also frequently observed (Galán-Olleros et al., 2024). Orthopedic problems, compounded by feeding dysfunction, lead to low weight, height, BMI, and micronutrient deficiencies (Motil et al., 2012). MDS shares with RTT overlapping yet distinct GI and musculoskeletal alterations. Feeding difficulties in males with MDS often begin in infancy, leading to poor weight gain and failure to thrive. Gastroesophageal reflux, chronic constipation, vomiting, and bowel obstruction from severe motility disturbances are reported. Musculoskeletal problems include early hypotonia, later spasticity, joint laxity, contractures, scoliosis, and fractures. These contribute to impaired growth, mobility, and quality of life in males with MDS, underscoring the need for therapies addressing both epigenetic and structural dysfunction (Allison et al., 2024; Pehlivan et al., 2024; Ta et al., 2022; Van Esch, 2020).

*MECP2* expression is not restricted to the nervous system. It is also expressed in peripheral tissues, including skeletal and visceral muscle (Pelka et al., 2005; Wang et al., 2021). While most RTT animal model studies have focused on the nervous system, it has been observed in *Mecp2* knockout mice an alteration of muscle architecture with hypotrophic fibers and tissue fibrosis (Conti et al., 2015). However, a muscle-specific *Mecp2* knockout mouse revealed no muscle defects, suggesting that the muscle and GI manifestations might be non-cell autonomous (Conti et al., 2015). A subsequent study in which loss of *Mecp2* has been limited to the peripheral tissues has reported hypoactivity and exercise fatigue (Ross et al., 2016). In addition, analysis of cardiomyocytes derived from *Mecp2* knockout embryonic stem cells revealed cell intrinsic alteration of development and function (Hara et al., 2015). Moreover, although multiple *Mecp2* overexpression and duplication mouse models faithfully reproduce the neurological features of MDS, including seizures, spasticity and premature death, detailed analyses of skeletal or visceral muscle structure and function have not been reported, leaving the contribution of excess MeCP2 in muscles essentially unexplored *in vivo* (Collins & Neul, 2022; Ezeonwuka & Rastegar, 2014). Overall, these studies returned an incomplete picture of the non-neuronal and potentially muscle-specific effects of alteration of *MECP2* expression, which might arise independently from defects in the nervous system and might not be fully recapitulated in existing mouse models (Vashi & Justice, 2019).

Here, we study the impact of human *MECP2* misexpression on muscle tissue unsing *Drosophila melanogaster*. This prime genetic model allows fine control of tissue- and time-specific *MECP2* expression, enabling precise assessment of its impact on muscle development and function. Although flies lack a *MECP2* ortholog, they have been extensively used to model the neurotoxic effect of human *MECP2* overexpression (Chen et al., 2025; Cukier et al., 2008; Sonn et al., 2025; Vonhoff et al., 2012; Williams et al., 2016). We report that *MECP2* misexpression in the *Drosophila* mesoderm is sufficient to cause skeletal and visceral muscle dysfunction, with effects that are strongest when misexpression occurs during early development. Interestingly, supplementation of histone deacetylase (HDAC) inhibitors ameliorates muscle structure and function in animal expressing *MECP2*, supporting the adoption of pharmacologic strategies directed at the modulation of chromatin accessibility for MECP2-related peripheral pathology.

## Methods

### Study design

We used *Drosophila melanogaster* to test whether human *MECP2* misexpression in mesodermal tissues causes skeletal/visceral defects, to define developmental windows of vulnerability, and to evaluate the ability of HDAC inhibitors to rescue defects. Primary outcomes were: (i) skeletal muscle morphology (larval muscles 6/7 and nuclear positioning), (ii) larval locomotion and adult negative geotaxis, and (iii) gut peristalsis and GI clearance. Secondary outcomes included neuromuscular junction (NMJ) morphology and mitochondrial ultrastructure (TEM). All experiments included genetic controls, ≥3 biological replicates. All data are presented as mean ± standard deviation, unless otherwise noted. For comparisons statistical significance was assessed using two-tailed unpaired t-tests. Analyses were performed in GraphPad Prism 8, with significance intervals denoted as ns p> 0.05, * p < 0.05, ** p < 0.01, *** p < 0.001, **** p < 0.0001.

### *Drosophila* husbandry and genetics

All strains were maintained and crossed at 25 °C, on standard diet (SD) composed of 93 grams of corn flour (Biosigma #78914), 25 grams of dried yeast (Biosigma #789126), 155 grams of molasses (Biosigma #789214), 7 grams of agar (Biosigma #789148), 70 ml of propionic acid (Carlo Erba #400-553) and 20 ml of 10% tegosept in EtOH (Biosigma #789130) per liter. Tissue-specific overexpression of human *MECP2* was achieved using the GAL4-UAS system (Brand & Perrimon, 1993). For temporal control, GAL80ts was employed: at 18 °C, ubiquitous expression of *tub-GAL80ts* represses GAL4 activity, whereas at 29 °C GAL80ts is inactivated allowing GAL4-dependent expression. The following fly lines were used: *Mef2-GAL4* (Jafar-Nejad Lab, Baylor College of Medicine, Houston, TX, USA (BCM), *how24B-GAL4* (Jafar-Nejad Lab, BCM), *vm-GAL4* (BDSC #48547), *c179-GAL4* (BDSC #6450), *hand-GAL4* (BDSC #66795), *tey-GAL4* (BDSC #2702), *elav-GAL4* (BDSC #8765), *gmr-GAL4* (BDSC #47463), *D42-GAL4* (BDSC #8816), *Ilp7-GAL4* (Miguel-Aliaga Lab, Francis Crick Inst.), *Mip-GAL4* (BDSC #51983), *NP3270-GAL4* (DGRC #113192), *NP3270-GAL4* (DGRC #113157), *mex-GAL4* (BDSC #91368), *tub-GAL80ts* (BDSC #7108); *UAS-GFP* (BDSC #5431), *UAS-nuclear LacZ* (Francis Crick Inst.), *UAS-MECP2* (Botas Lab, BCM), *UAS-MECP2^R106W^* (Botas Lab, BCM), *UAS-MECP2^R294X^* (Botas Lab, BCM). Genotypes of all samples in the study are listed in Table S1.

### Developmental Progression, Survival, and Fitness Assays

Collections of eggs laid over 24 hours were incubated at 25 °C on control or supplemented food. To quantify global developmental progression, the proportion of animals at each developmental stage (larvae, early pupae, late pupae, adults) was assessed at 12 days AEL. All individuals in a tube were scored and represented in a stacked-bar plot.

To assess lethality, the GAL4 drivers used for *MECP2* misexpression were grouped based on their predominant expression domains (ubiquitous, nervous system, enteric neurons, mesoderm, and enterocytes) and survival was calculated relative to expected Mendelian ratio. Tissue-specific patterns were tested by crossing each driver to *UAS-GFP* and evaluating the GFP signal in L3 larvae by immunofluorescence.

### Immunofluorescence analyses

For body wall muscles and NMJ immunolabeling, third-instar larvae (L3) were dissected in PBS and fixed in 4% PFA (20 min). Samples were washed in 0.1% PBST, blocked in 3% BSA, and incubated overnight at 4 °C with: Anti-MECP2 (Invitrogen #PA1-887; 1:200), Mouse anti-Bruchpilot (nc82; DSHB; 1:100). After washes, tissues were stained 2 hrs at room temperature with: Alexa-488 anti-rabbit (Jackson #11-605-003; 1:200), Cy3-anti-HRP (Jackson 123-165-021; 1:200), Phalloidin-TRITC (Sigma-Aldrich #P1951; 1:1000). Nuclei were counterstained with DAPI (Merck Life Sciences #D9542; 1:5000). For visceral muscles immunolabeling, larval and adult guts were fixed in 4% PFA (40 min), washed twice in PBS and stained for 1.5 hrs at room temperature with Phalloidin-AlexaFluor 647 (Thermo Fisher Scientific #A22287; 1:500), mounted in Vectashield with DAPI (Vector #CA 94010) as described in Mineo *et al*., 2024. For counterstaining with anti-MECP2, the protocol was modified with the permeabilization, blocking, overnight primary antibody incubation, and secondary antibody staining steps used for body wall muscles. To evaluate *Mef2* vs *c179* embryonic expression, embryos of the appropriate stage were collected on apple juice agar plates, treated briefly with 100% bleach for 1–2 min to remove the chorion, and washed thoroughly with distilled water. Embryos were then dried on filter paper and transferred to clean tubes. Fixation was performed in a scintillation vial containing heptane, PBS, and 4% formaldehyde for 20 min with constant shaking. After removal of the aqueous layer, methanol was added, embryos were vigorously shaken to remove the vitelline membrane and accumulated in methanol (Müller, 2008). Embryos were then washed at least twice and stained with anti-GFP 1:1000 (Abcam #ab5450) to detect transgene expression, anti-FasIII 1:50 (DSHB #7G10) to visualize visceral muscle membranes, and DAPI (Vectashield + DAPI, ThermoFisher #NC9524612) to visualize the nuclei.

For quantifications, confocal images, acquired with a Leica SP8 confocal microscope, were analysed in Fiji (Schindelin et al., 2012). For the calculation of the muscle fiber width/length ratio, measurements were taken at the center of hemisegments A2-A4. For each larva, the mean width/length ratio was calculated for both the right and left muscles 6 and 7, and this average was plotted as a single data point for each muscle. For nuclear analyses, nuclei from muscles 6 and 7 in hemisegments A2–A4 were quantified using confocal images stained with Phalloidin and DAPI. Nuclear parameters included: nuclear area; nuclear circularity (defined as length/width, with values closer to 1 indicating more circular nuclei); nuclear solidity (solidity ratio is defined as ratio of a nucleus area to the area of its convex hull); and nuclear positioning, measured as the average internuclear distance. For morphological analysis of larval muscles 6 and 7, individual branches were counted manually along the main axonal arborization, discrete synaptic boutons were identified using anti-HRP staining and counted manually. Puncta marked by anti-Bruchpilot were quantified within the area defined by the anti-HRP signal, with integrated density (IntDen) measured in FIJI after thresholding each NMJ.

### Transmission Electron Microscopy

Body-wall muscle specimens were fixed in 2.5% glutaraldehyde prepared in 0.1 M sodium cacodylate buffer (pH 7.4) for 3 h at room temperature, followed by extensive buffer rinses. Samples were post-fixed in 2% osmium tetroxide in the same buffer for 2 h, rinsed in distilled water, and contrasted *en bloc* with 1% aqueous uranyl acetate for 1 h. Following fixation, samples were dehydrated through a graded ethanol series and embedded in epoxy resin according to standard protocols. Ultrathin sections (∼50 nm) were obtained using a Leica UCT ultramicrotome, collected on copper grids, and counterstained with uranyl acetate prior to imaging. Sections were examined using a Hitachi HT7800 transmission electron microscope operating at 120 kV (Hitachi, Tokyo, Japan). Digital micrographs were acquired with MegaviewG3 camera at comparable magnifications for all experimental conditions. Mitochondrial number and cristae morphology were quantified using Radius v2.0 software. For each condition, multiple cells and sections were analyzed, and normal versus aberrant mitochondria were identified based on established ultrastructural criteria (Bellese et al. 2023).

### Behavioral and Functional Assays

To assess larval locomotion, individual L3 larvae were placed on a wet 0.5 cm-grid and allowed to crawl for 60 seconds. Distance traveled was calculated as number of gridlines crossed × 0.5 cm. To evaluate L3 larvae wandering ability, the amount of food in each vial and the egg-laying time was controlled to ensure equal larval density. Wandering ability was quantified by measuring the linear distance between the food surface and the final position of the pupal case on the vial wall.

To assess feeding behavior, a comparable number of L3 wandering larvae were collected from eggs laid by equal numbers of parental flies. At approximately 96 hours after eggs laying (AEL), early wandering L3 larvae were gently harvested under a stereomicroscope using fine-tip forceps and transferred in groups of five into silicone-coated Petri dishes containing a thin layer (∼2 mm) of 2% yeast solution. Following a 30-second acclimation period, larval movements were video-recorded at 10× magnification for 60 seconds. Individual mouth hook contractions were subsequently counted from the recordings, and the mean contraction rate per minute was calculated for each group.

To quantify climbing ability, females were tapped to the vial bottom and given 10 s to climb above a 7 cm mark. Trials were repeated four times per group at 5, 10, 20, and 30 days post eclosion (dpe); the climbing index is the mean proportion above the mark. In case of supplementations, animals were fed supplemented diets since hatching.

To assess the muscle-shaking phenotype, adults expressing *MECP2* under the *c179-GAL4* muscle driver were evaluated at 30 dpe. The phenotype was defined as a compulsive, abrupt, and uncoordinated displacement of flies from one point of the vial to another upon mechanical stimulation. For each assay, the fraction of animals displaying shaking behavior was averaged to obtain a single value per measurement.

To study gut peristalsis, intact guts of early L3 larvae were dissected in Schneider’s medium (Thermofisher #21720024) and peristaltic contractions counted for 1 min under a stereomicroscope. To evaluate the gut clearance, wandering L3 larvae fed on diet supplemented with 0.005% bromophenol blue (BPB; Merck #B8026) were placed on Whatman paper wetted with 20% sucrose/PBS. Clearance was quantified as the percentage of animals that had retained food in their gut, calculated as blue-positive larvae / total larvae at different time after picking (TAP).

### Dietary Supplementation

NaB (Sigma-Aldrich #B5887), AcOH (Sigma-Aldrich #27221; Martelli et al., 2024), Lalbaay® (Kolfarma SRL #982845036; NCT05420805) and VPA (Merck #P4543) were added to freshly cooked SD cooled to 65°C. Final concentrations in the food were: 20 mM NaB, added from a 10x stock solution (Di Fede et al., 2021); 0.75% AcOH (Martelli et al., 2024), Lalbaay®, adjusted to obtain final concentration of 20 mM NaB, and 1 mM VPA (Di Fede et al., 2021). Flies were reared on supplemented or control medium from hatching to adulthood. In supplementation experiments, survival was expressed as the observed vs. expected ratio of progeny. For normalization, the average survival of the control genotype on standard diet was set to 100%. To estimate pupal survival, larvae were reared on standard diet or diets supplemented and survival was calculated as the percentage of viable pupae over the total number of pupae scored. Pupae displaying a rigid, elongated cuticle and failing to progress to pharate adult stage were scored as non-viable. Developmental delay was quantified as the ratio of pupae to larvae at 8 days AEL, using progeny from synchronized parentals. For the gut clearance assay with supplements, the food retention index was calculated as (No discharge×1 + Semi discharge×0.5 + Full discharge×0) / (Total larvae scored), and the residual gut content was scored at different times after picking. To evaluate supplement toxicity, the survival ratio - calculated as previously described - was measured under the different treatments and compared with SD.

## Results

### Mesoderm-specific *MECP2* overexpression causes lethality during development

To determine the impact of human MeCP2 on *Drosophila* tissues, we first assessed survival following misexpression driven by a panel of GAL4 drivers. Mesodermal drivers (*Mef2-GAL4*, *how24B-GAL4*, *c179-GAL4*, *vm-GAL4* and *cg-GAL4*) showed the strongest effects, with reduced or complete absence of adults at 12 days AEL, thus recapitulating the lethality observed by ubiquitous expression (*actin-GAL4*; Table S2). While expression with *how24B-GAL4*, resulted in lethality at the late pupal stage, expression with *Mef2-GAL4* causing lethality in white pupae (Figure S1a), consistent with the fact that *Mef2-GAL4* drives expression in the mesoderm earlier than *how24B-GAL4* (Galeone et al., 2017). Both *Mef2>MECP2* and *how24B>MECP2* animals progressed through larval development and underwent pupariation, with pupae that were elongated compared to control. In addition, some of *Mef2>MECP2* larvae did not fully pupariate (Figure S1b–e’). Mouth hook contraction frequency, a direct readout of feeding activity, was indistinguishable between *Mef2>MECP2* and control larvae (Figure S1f; Video S1), indicating that such phenotypes are not due to reduced food intake. Together, these data strongly suggest that the mesoderm is highly sensitive to *MECP2*-induced perturbations.

### *MECP2* overexpression impairs muscle morphology and function in *Drosophila* larvae

Considering that *MECP2* misexpression alters development, we evaluated the morphology and functionality of larval muscles, which are derived from the mesoderm. We first assessed whether MeCP2 can be found in *Drosophila* muscles. Consistent with the subcellular localization previously reported in neurons (Cukier et al., 2008), immunostaining to detect MeCP2 localization revealed nuclear localization in skeletal body wall muscles and visceral gut muscles of *Mef2>MECP2* larvae (Figure S2), suggesting that the human protein might be active in mesoderm-derived tissue. Then, we examined the morphology of muscle 6 and 7 in segment A2-A4 in *Mef2>MECP2* and control larvae (Figure 1a). Compared to *Mef2>GFP* controls, animals expressing *MECP2* showed prominent thinning of the abdominal body wall muscle, (Figure 1b, quantified in b’). In addition, *MECP2* expression caused elongation, deformation and mispositioning of nuclei within fibers (Figure 1c quantified in c’). We also analyzed the ultrastructure of larval body wall muscles and found that *MECP2*-expressing larvae have more mitochondria than controls and these often possess disorganized cristae (Figure 1d, quantified in d’). Then, we determined whether these morphological changes were accompanied by functional impairments. Larval locomotion and wandering ability, two readouts of skeletal muscle function, were significantly reduced in *MECP2*-expressing larvae, when compared to controls (Figure 1e-f). These defects do not correlate with alteration of NMJs (Figure S3), suggesting that they are not induced by changes in the ability of muscles to receive appropriate stimulation. These data reveal that muscle morphology and function are cell-autonomously altered by MeCP2.

**Figure 1.**
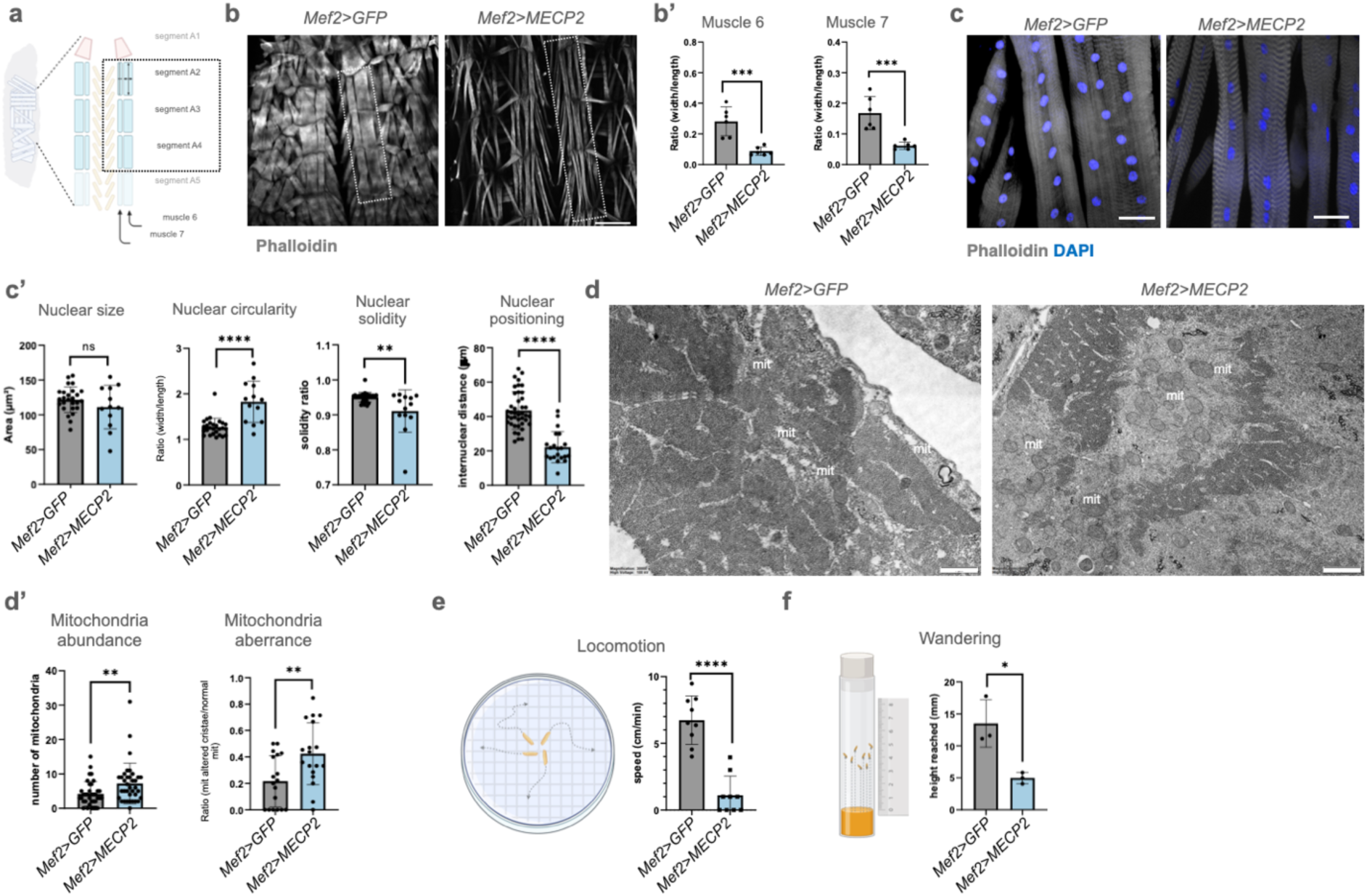
mesodermal *MECP2* misexpression impairs skeletal muscle morphology, mitochondrial integrity and motor function. **(a)** Schematic representation of the central region of the body wall muscles of an L3 larva. The reference abdominal hemisegments A2-A4 of muscles 6 and 7 that we analyzed are boxed. **(b)** Phalloidin staining of abdominal body wall muscles of the indicated genotypes. The box indicates the quantified region. Scale bars: 150 µm. **(b’)** Quantification of muscle 6 and 7 width/length ratio as described in methods (N = 6 larvae). **(c)** Representative confocal images of abdominal muscles 6 and 7 stained with phalloidin (white) and DAPI (blue). **(c’)** Quantification of nuclear parameters as described in the methods. N > 3 biological replicates, each dot represent an eligible nucleus. **(d)** Transmission electron micrographs of L3 body-wall muscle from *Mef2>GFP* controls and *Mef2>MECP2* animals. mit Mitochondria. Scale bars, 1 µm. **(d’)** Quantification of mitochondrial number per muscle section and the % of mitochondria with altered cristae. Morphometric analysis was performed on 20 aligned TEM micrographs per genotype (20k magnification) from 3 biologically independent samples. **(e)** Schematic illustration of the quantification of the larval speed, which was calculated as described in methods. Each point represents a single larva (N = 9 larvae per sample). **(f)** Schematic illustration of the quantification of the height reached by L3 larvae prior to pupariation. Each point represents the average of 30 pupal cases per biological replicate (3 biologically independent replicates).

We next asked whether larval visceral muscles were affected by *MECP2* expression (Figure 2a). Interestingly, we observed that the GI tract of *Mef2>MECP2* animals is shorter than that of control animals (Figure 2b, quantified in b’). In addition, direct analysis of the gut muscles revealed ruptures of circular muscle fibers in ∼60% of *Mef2>MECP2* animals (Figure 2c). We next assessed food transit and peristalsis. To measure food transit, we monitored discharging of the bromophenol blue-containing fly food developmentally occurring at the L3 wandering stage. In contrast to *Mef2>GFP* larvae that all discharge food by 24 h after picking, ∼50% of *Mef2>MECP2* larvae retained colored food, in some cases retaining it after pupal cuticle formation (Figure 2d, quantified in d’). A milder phenotype was observed with the later onset *how24B-GAL4* driver (Figure S4). Consistent with this, the peristaltic activity in the dissected GI of *Mef2>MECP2* larvae was strongly reduced compared to control, with significantly fewer contractions per minute (Figure 2e, Video S2). Together, these data show that *MECP2* misexpression impairs visceral muscle activity, resulting in defective GI tract motility and food retention.

**Figure 2.**
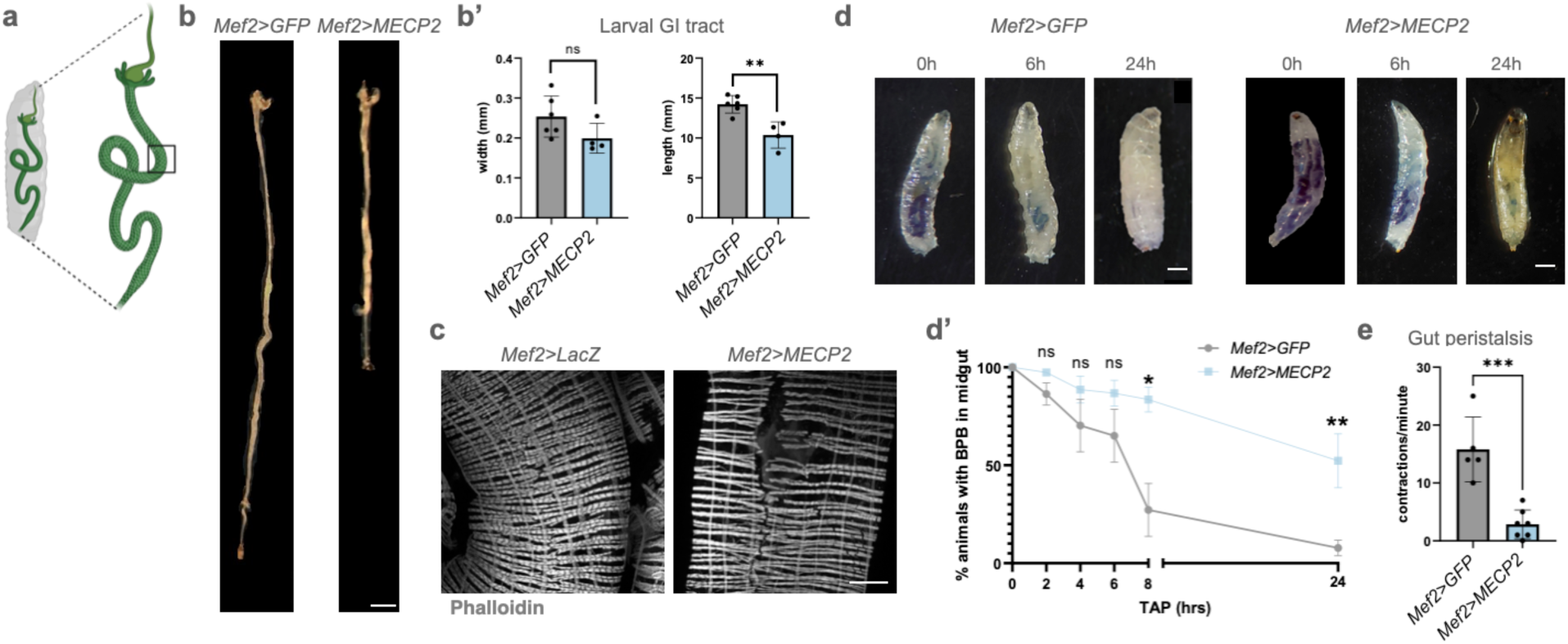
*MECP2* overexpression in muscle impairs GI morphology and function in *Drosophila* larvae. **(a)** Schematic representation of a L3 larva with the GI tract highlighted and the dissected tract showing longitudinal and circular visceral muscles. The black box indicates the region of the midgut where high magnification confocal images were acquired. **(b)** Representative bright field images of the larval GI tract from control *Mef2>GFP* and *Mef2>MECP2* animals. Scalebar 1mm. (**b’)** Quantification of GI length and width. Points represent individual animals; (N≥4 animals). **(c)** Representative confocal microscopy images of a midgut tract labeled with phalloidin. Approximately, 60% of *Mef2>MECP2* larvae show fiber ruptures of the circular muscles (N=10 larvae). **(d)** Representative bright-field images of L3 larvae. Control *Mef2>GFP* and *Mef2>MECP2* larvae were fed blue food and imaged at the indicated time after picking to monitor gut content. Scale bar: 1 mm. **(d’)** Graph representation of the percentage of larvae that retain dyed food over time quantified as described in methods. Control larvae efficiently cleared the food 24 hrs after picking, indicating normal GI transit. In contrast, ∼50% of *Mef2>MECP2* larvae retained the colored food, in some cases beyond pupal cuticle formation. N ≥ 30 larvae per genotype, ≥3 independent experiments. Data are mean ± SEM. **(e)** Bar graph of peristaltic contractions per minute in dissected GI preparations. Each point represents one larva (N≥5 larvae).

### Spatiotemporal *MECP2-*induced perturbation of adult muscles

To understand when MeCP2 most affects gut muscle development, we used *tub*-*GAL80ts* to block *Mef2-GAL4* expression during early development (*Mef2ts>* hereafter; see Methods). At 18°C, this manipulation prevented early lethality and allowed morphologic analysis of the adult gut (Figure 3a). We induced expression by shifting crosses at 29° C immediately after eclosion (Figure 3b), or after gut maturation at 5 dpe, and analyzed animals at 10 dpe (Figure 3c). Upon induction at eclosion, adult guts from *Mef2ts>MECP2* flies exhibited altered morphology, with increased width and disorganized muscle fibers, when compared to controls (Figure 3d-e). In contrast, induction after gut maturation produced only a mild shortening of the midgut and more subtle changes in fiber organization (Figure 3f-g). To test whether the gut muscle phenotypes can be achieved by limiting *MECP2* expression to a developing set of circular midgut muscles (Buchon et al., 2013; Kuroda et al., 2012) rather than the whole mesoderm, we used *Hand-GAL4.* Consistent with our previous evidence, the midgut of 40% of the *Hand>MECP2* adults displayed morphologic alteration of the gut muscles as well as shortening, together with extremely reduced peristaltic activity (Figure 3h-j). Similar alteration in gut length, width or morphology were also observed when *MECP2* was expressed specifically in longitudinal gut muscles (Klapper et al., 2001) using the *tey-GAL4* driver or with the alternative visceral muscle driver *vmTS-GAL4* (Figure S5). Together with our previous data, these results indicate that *MECP2* expression predominantly impacts muscles during embryonic, larval and early adult development, with limited effects after gut maturation, and that expression in visceral muscle is sufficient to compromise gut muscle morphology and function.

**Figure 3.**
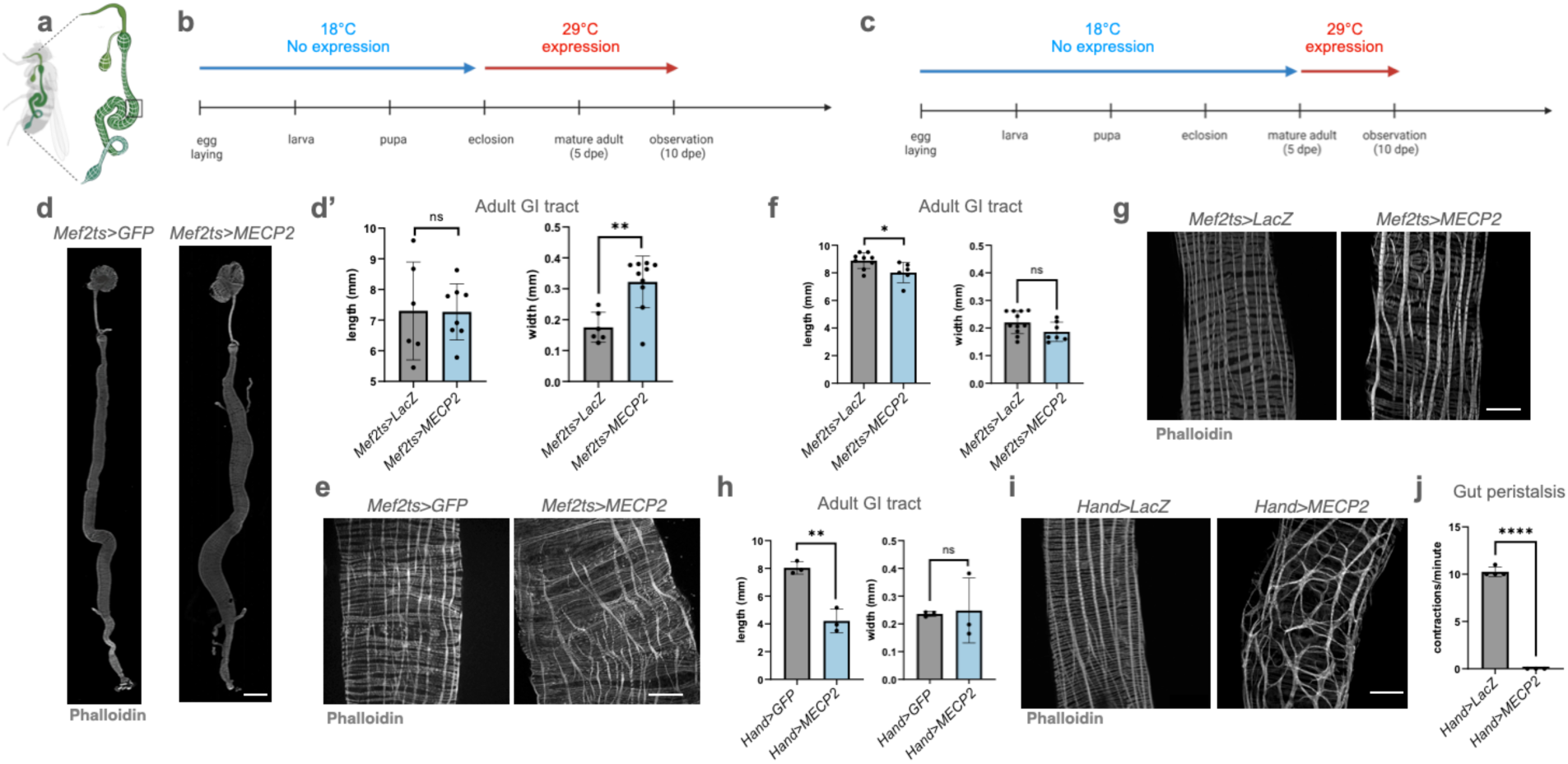
**(a)** Schematic representation of an adult fly with the GI tract highlighted and the dissected tract showing longitudinal and circular visceral muscles. The black box indicates the region of the midgut where high magnification confocal images were acquired. **(b)** Illustration of the timeline used to control *MECP2* expression under the *Mef2*-*GAL4 GAL80ts* driver combination. The temperature-sensitive GAL80ts repressor blocks GAL4 activity at 18 °C and allows *MECP2* expression at 29 °C immediately after eclosion until observation time at 10 dpe (early induction). **(c)** Illustration of *MECP2* expression at 5 dpe, once visceral muscle development is complete, until observation time at 10 dpe (late induction). **(d)** Confocal images of representative GI tracts from adult flies with early induction of *MECP2* expression. Scalebar: 500μm. **(d’)** Quantification of gut length and width under early induction (N=5-10 GI tract per sample). **(e)** Confocal images of representative phalloidin-stained visceral muscles of the midgut upon early induction of MECP2 expression (N=10 per sample analyzed). Scalebar: 50μm. **(f)** Quantification of gut length and width in flies with late induction of *MECP2* expression (N>8 GI tract per sample). **(g)** Representative confocal images of phalloidin-stained midguts under late induction (N=15 per sample analyzed). Scalebar: 50μm. **(h)** Quantification of gut length and width in *Hand>MECP2* and control adults. Each dot represents the mean of 3 animals. (**i)** Confocal images of adult midguts stained with phalloidin in *Hand>MECP2* and control animals (N=15 per sample analyzed). **(j)** Number of peristaltic contractions per minute in adult guts of the indicated genotypes. Each point represents a gut.

To assess performance of the adult skeletal muscles, we used the *c179-GAL4* driver whose expression begins at the larval stage, rather than during embryogenesis (as is the case of *Mef2-GAL4*; Figure S6a-b). Compared to control, *c179>MECP2* adults display a pronounced age-dependent progressive impairment in climbing ability and evident muscle shaking (Figure S6c-d, Video S3), indicating that adult skeletal muscle function can also be compromised by *MECP2* misexpression.

### Developmental NaB supplementation partially rescues the muscle alterations of *MECP2*-expressing flies

Considering the role of SCFAs as HDAC inhibitors (Kabir et al., 2025; Kasubuchi et al., 2015), we tested whether dietary supplementation from larval hatching could modulate the phenotypes observed upon *MECP2* expression (Figure 4a). We first quantified adult survival of *c179>MECP2* flies, which was found very limited in SD (Figure 4b). Interestingly, we found that NaB supplementation or supplementation with Lalbaay® (Firoozi et al., 2024; Guan et al., 2021) containing equimolar NaB concentration, or AcOH supplementation, all increased adult survival of *c179>MECP2* animals, with NaB and Lalbaay® providing the strongest increase (Figure 4b). NaB and Lalbaay® supplementation improved also the larval wandering ability of *Mef2>MECP2* larvae (Figure S7a) and was not toxic, while AcOH was less tolerated (Figure S7b). Because of this and of the fact that Lalbaay® contains NaB, we next focused on the effect of NaB. Supplementation of NaB from larval hatching also ameliorated the climbing ability of *c179>MECP2* adults at 10 dpe (Figure 4c), as well as it prevented the emergence of marked defects in larval skeletal muscles (Figure 4d, quantified in d’), and partially relieved the GI transit impairment and the limited gut peristalsis observed in *Mef2>MECP2* larvae (Figure 4e-f).

**Figure 4.**
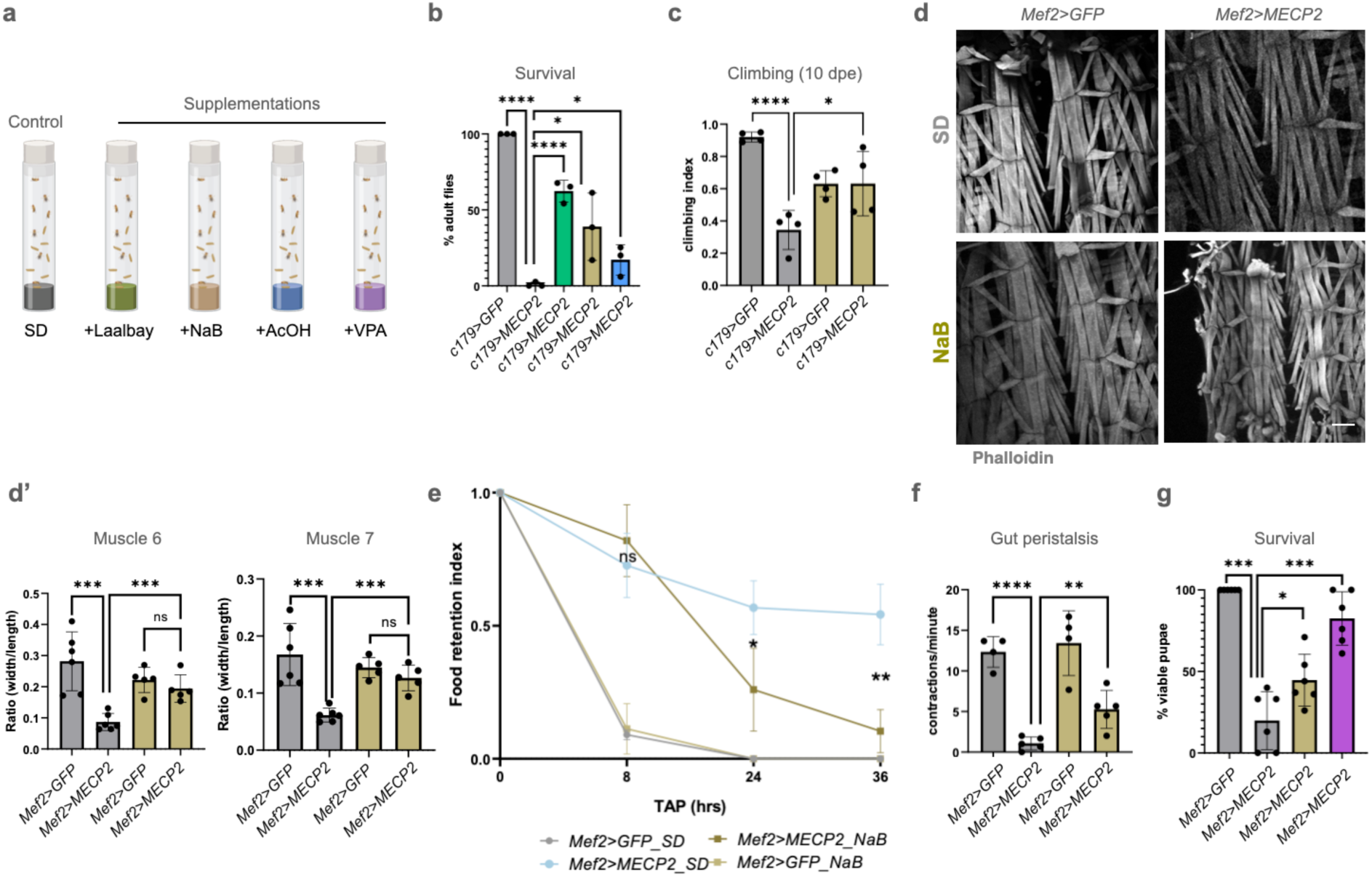
SCFA supplementations: beneficial effects of butyrate in *MECP2*-overexpressing flies. **(a)** The schematic illustrates the standard diet (SD, shown in grey) and its supplementation with different compounds (shown in color, see Methods). **(b)** Bar graph showing adult survival from 3 independent biological replicates under different dietary conditions (N=80 animals per replicate). **(c)** Climbing assay at 10 dpe. Each point represents the mean of 3 technical measurements, each performed on 10 flies (N=4 biological replicates). **(d)** Confocal images of larval body wall muscles stained with Phalloidin. Scale bar: 150 μm. **(d’)** Quantification of the width/length ratio of body wall muscles 6 and 7 of hemisegment A2-A4 (N=5 larvae). **(e)** Quantification of the food retention in larvae of the indicated genotype fed as indicated as described in methods. Mean ± SEM are shown for 5 independent biological replicates (N = 10 larvae per replicate). **(f)** Quantification of peristaltic contractions per minute in larvae of the indicated genotypes. The average of 3 animals per biological replicate (N=5) was used. **(g)** Bar graph showing pupal survival of *Mef2>GFP* and *Mef2>MECP2* larvae from 5 biological replicates, expressed as the percentage of viable pupae over the total number of pupae. N=80 per replicate. Larvae were supplemented as indicated.

To distinguish between the potential trophic effect of SCFAs and their effect as a HDAC inhibitor, we supplemented the SD with VPA, a broad-spectrum HDAC inhibitor lacking nutritional properties (Di Fede et al., 2021). Remarkably, VPA recapitulated the beneficial effects of NaB, improving both development progression (Figure S7c) and survival (Figure 4g) of *Mef2>MECP2* animals. Overall, these findings support the possibility that NaB exerts its beneficial effects via regulation of altered deacetylation.

### *MECP2* mutants link muscle phenotypes to RTT pathogenesis

To determine whether the observed mesodermal-induced phenotypes are relevant to MeCP2-dependent diseases, we compared misexpression of MeCP2 with that of 2 MeCP2 variants typically found in RTT patients (Figure 5a). MeCP2^R106W^ originates from a missense mutation in the region encoding the highly conserved methyl-CpG binding domain (MBD) and is consistently associated with severe clinical outcomes (Mietto et al., 2025; Wan et al., 1999). MeCP2^R294X^ is characterized by a nonsense mutation that results in protein lacking the transcriptional repression domain (TRD) (Ananiev et al., 2011; Chae et al., 2002) and is generally causing mild disease presentations (Cuddapah et al., 2014; Jara-Ettinger et al., 2021). Interestingly, compared to *Mef2>MECP2*, *Mef2>MECP2^R106W^*animals survived to adulthood. In sheer contrast, *Mef2>MECP2^R294X^*animals survived less than *Mef2>MECP2* animals (Figure 5b). Consistent with this, pupae expressing *MECP2^R106W^* were of normal size, while those expressing *MECP2^R294X^* failed to pupariate and elongated as *MECP2*-expressing larvae (Figure 5c, quantified in c’).

**Figure 5.**
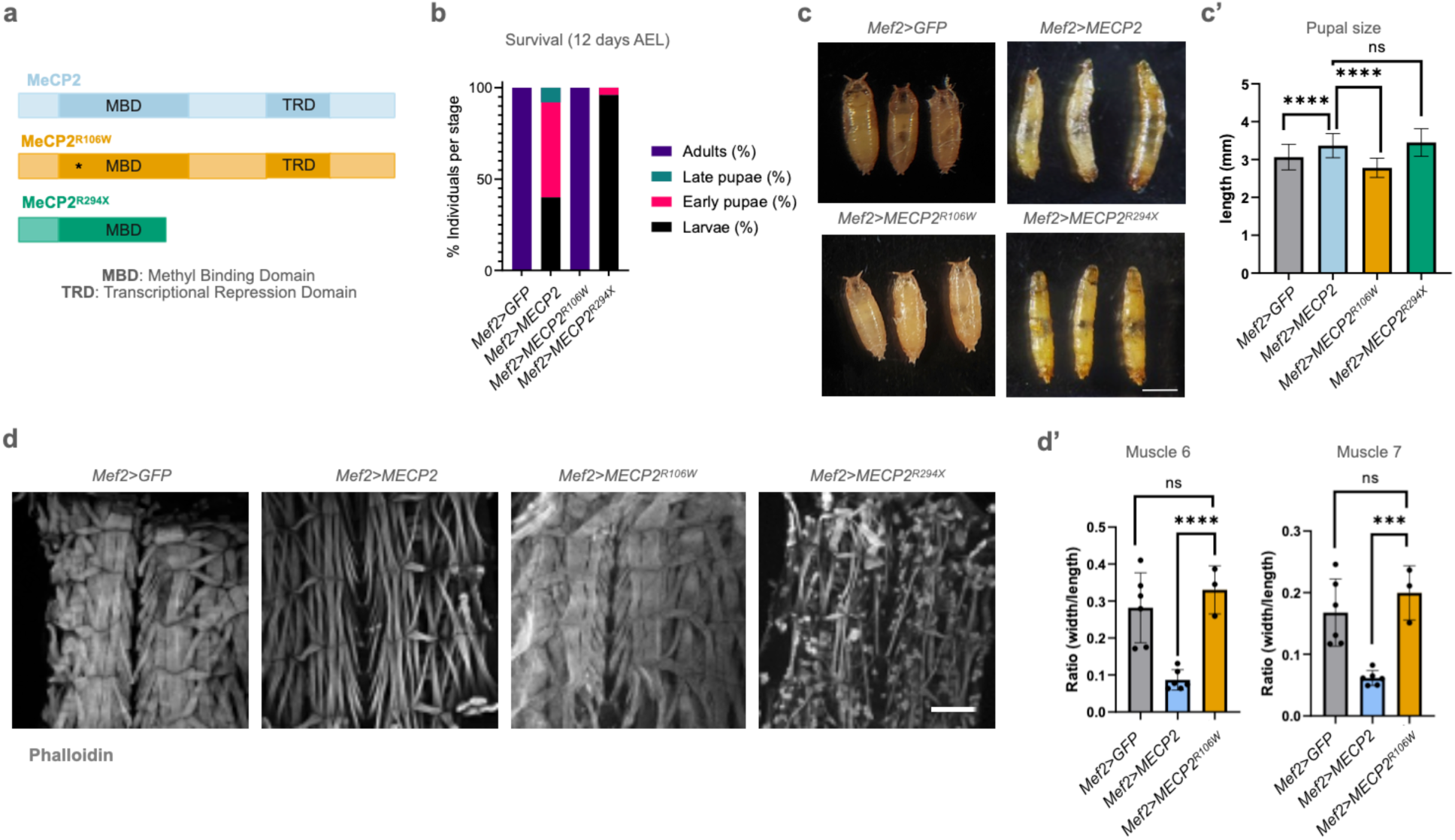
*MECP2* variants misexpression in the fly mesoderm as a genetic measure of RTT variant pathogenicity. **(a)** Schematic representation of MeCP2 variants with indication of MBD (methyl-CpG binding domain,) and TRD (transcriptional repression domain) domain boundaries. The asterisk indicates the position of the missense mutation causing the R106W aa change. **(b)** Stacked bar chart showing the % of animals at each of the indicated developmental stages when fly populations are analyzed at 12 days AEL for the indicated genotype (N=30 animals). **(c)** Representative images of 3 early pupae of the indicated genotype. Scale bar: 1 mm. **(c’)** Quantification of pupal length for the indicated genotypes. (N=50 animals) **(d)** Confocal images of larval body wall muscles stained with phalloidin. Scale bar: 150 μm. **(d’)** Quantification of the width/length ratio of body wall muscle 6 and 7 in hemisegments A2-A4 plotted for the indicated genotypes (N≥3 biological replicates). Quantification was not performed for MeCP2^R294X^ because the phenotype was too severe to allow consistent measurement.

Visualization of body wall muscles revealed that MeCP2^R106W^ did not induce muscle thinning while MeCP2^R294X^ induced strong thinning and degeneration (Figure 5d quantified in d’). Thus, our genetic analysis indicates that gain-of-function phenotypes of mesodermal MeCP2 expression are not recapitulated by the loss-of-function MeCP2^R106W^ variant, while the partially function-retaining variant MeCP2^R294X^ yields gain-of-function phenotypes that are even stronger that of MeCP2.

## Discussion

Although *Drosophila* does not possess a *MECP2* ortholog and shows minimal levels of methylation on its genomic DNA (Capuano et al., 2014), ectopic expression of human *MECP2* in the *Drosophila* nervous system recapitulates neurotoxicity, chromatin association, phosphorylation at serine 423 and genetic interaction with *Drosophila* orthologs of known MeCP2 partners (Cukier et al., 2008). Now, we show that restricting human *MECP2* expression to the *Drosophila* mesoderm is sufficient to disrupt development and function of both skeletal and visceral muscles, leading to impaired motor function, gut dysmotility, which are often observed in patients affected by MeCP2 diseases.

While mesodermal overexpression of *MECP2* might more directly model MDS, the extent to which our findings are also informative for RTT, is less obvious. Recent comparative expression studies have used *Drosophila* genetic to functionally stratify MeCP2 variants associated with autism and RTT (Chen et al., 2025). In line with this approach, we find that the MeCP2^R106W^, which show weakened MBD binding to DNA and is associated with severe RTT, behaves as a functional loss-of-function variant and failed to reproduce the toxic effects of MeCP2 in muscles, whereas the truncated variant MeCP2^R294X^, lacking the TRD domain, exacerbated developmental arrest and muscle defects, in agreement with previous findings in the fly nervous system. Thus, our data predicts that the pathogenic mechanisms at play in central and peripheral tissues might be similar and that our assays might also capture clinically relevant aspects of RTT pathology. Future work will point to testing more disease-associated MeCP2 variants and will clarify the nature of the different gain-of-function effects of MeCP2 and MeCP2^R294X^.

Non-neuronal contributions to MECP2-related traits remain incompletely defined. In mice, skeletal muscle–specific deletion of *Mecp2* did not produce overt structural muscle defects (Conti et al., 2015), whereas re-expression of Mecp2 in the nervous system in a null background revealed abnormalities in several mesoderm-derived organs, including bone and kidney, and caused reduced body weight, hypoactivity, and exercise fatigue, but the gastrointestinal tract was not systematically examined (Ross et al., 2016). Together with the low endogenous MeCP2 levels reported in mouse skeletal muscle and many peripheral tissues, these findings have often been interpreted as arguing against a major muscle-autonomous component. However, immunohistochemical analyses and RNA-seq data from the Human Protein Atlas (Human Protein Atlas, accessed 2025; Ross et al., 2016; Shahbazian et al., 2002) indicate robust *MECP2* expression in skeletal and visceral muscle, suggesting that peripheral phenotypes may be more cell-autonomous in patients than in mice. Thus, our temporally and spatially restricted *Drosophila* models of peripheral MeCP2 might overcome some potential limitations of mouse models. Although we cannot study all human mesoderm-derived organs in flies (*i.e.* skeletal system), our analysis could extend to more mesodermal organs that are conserved in flies (*i.e.* kidney) and affected in MECP2 related diseases in the future (Ward et al., 2016).

Temporal control of *MECP2* expressions in flies revealed decreasing gut vulnerability as development progressed. This early sensitivity is consistent with the dosage dependence of chromatin regulators during lineage specification and matches clinical observations that hypotonia and gastrointestinal dysmotility manifest early in RTT and often persist throughout life (Baikie et al., 2014; Motil et al., 2012). These findings suggest that prenatal and early postnatal periods may be particularly sensitive to MeCP2 imbalance, with later interventions potentially stabilizing or partially reversing established defects but less able to prevent their initial emergence. In practical terms, the timing of both genetic and pharmacologic interventions should be considered a key design variable in preclinical studies.

Supplementation experiments indicate that SCFA and valproate can mitigate MeCP2-induced muscle and gut dysfunction. Notably, direct interaction between MeCP2 and HDAC-containing co-repressor complexes have been widely reported (Collins & Neul, 2022) and previous evidence indicated that HDAC inhibition can restore microtubule dynamics and reverse early exploratory and neurological deficits in *Mecp2* null mice (Lebrun et al., 2021; Guo et al., 2014; Ishiyama et al., 2019). In this context, our data predict that early treatment with supplements containing HDAC inhibitors, such as Lalbaay®, which has been recently tested in RTT patients (ClinicalTrials.gov Identifier: NCT05420805), might be effective also to reduce symptoms in peripheral tissues of RTT and MDS patients. Considering that VPA, a synthetic HDAC inhibitor and antiepileptic drug without nutritional value (Göttlicher et al., 2001; Löscher, 1999), phenocopies the beneficial effects of NaB supplementation, we conclude that SCFA effects are due to epigenetic regulation rather than a direct impact as nutrients.

In terms of muscle morphology, the thin appearance and the nuclear misplacement observed in MeCP2-containing muscles suggest an early arrest in muscle maturation, a phenotype previously linked to alteration of glycolytic genes (Graca et al., 2021). Because reports in *Mecp2* null mice and RTT patients and our models have highlighted mitochondrial dysfunction (Gold et al., 2014; Shulyakova et al., 2017), future transcriptional and chromatin profiling of tissue from SCFA supplemented versus unsupplemented MeCP2-expressing flies will reveal whether epigenetic regulation impact expression of genes related to nutrient metabolism.

## Conclusion and limitations of the study

Our work establishes that dysregulated MeCP2 activity confined to mesodermal tissues is sufficient to impair the development and function of both skeletal and visceral muscles. By defining an early developmental window of vulnerability and showing that SCFA and HDAC inhibition can partially normalize muscle architecture, locomotion, and gut motility, our *Drosophila* model provides a tractable *in vivo* platform to probe peripheral mechanisms of pathophysiology in MECP2-related disorders and to test interventions. At the same time, our conclusions are tempered by key limitations: flies lack a native MeCP2 ortholog and differ from mammalian muscle and GI physiology. Also, we rely on tissue-directed misexpression rather than physiological modulation of endogenous Mecp2 and we do not yet have direct chromatin or transcriptional readouts of HDAC inhibitor action. Our findings should also be viewed as a proof-of-concept warranting re-evaluation of vertebrate models of peripheral disease and could inform preclinical studies aimed at targeting HDACs and SCFA pathways to ameliorate disabling motor and gastrointestinal manifestations in RTT and MDS.

## Data sharing

*Drosophila* stocks, reagents and protocols, when not available at listed sources, can be requested to the corresponding author.

## Acknowledgements

T.V. acknowledges the support of PRIN grant 2022JKEBB8 and 2022PNRRYA3LL and of Telethon Italia. G.C. was a student of the PhD program in Translational Medicine of the University of Milan. I.M-A. is supported by MRC intramural funding and the Francis Crick Institute, which receives its core funding from Cancer Research UK (FC001317, FC001175), the UK Medical Research Council (FC001317, FC001175) and the Wellcome Trust (FC001317, FC001175). K.C. thanks the University of Genova for funding the acquisition of a Hitachi 120kV TEM HT7800 (grant DR3404, heavy equipment).

## Contributors

T.V. and A.G. designed the study and wrote the original draft. G.C. performed most of the experiments with the help of M.K.K. and A.M. M.C.G. and K.C. performed the EM experiments. A.V: and E.B. contributed to the formal analyses. I.M-A., E.B., A.G. and T.V. contributed to supervision, funding acquisition and to review and editing of the final manuscript.

