## Supplementary material for "Mesodermal-specific *MECP2* expression in *Drosophila* induces visceral and skeletal muscle defects rescued by butyrate supplementation": supp figures and videos

**Supplementary Data**

| Sample | Full Genotype |
|---|---|
| **Muscle drivers** | |
| *Mef2 > MECP2* | *yw; +; Mef2-GAL4/pUAS-MECP2* |
| *Mef2 > MECP2^R106W* | *yw; pUAS-MECP2-R106W/+; Mef2-GAL4/+* |
| *Mef2 > MECP2^R294X* | *yw; +; Mef2-GAL4/pUAS-MECP2-R294X* |
| *Mef2 > GFP* | *yw; pUAS-GFP/+; Mef2-GAL4/+* |
| *Mef2TS > LacZ* | *yw; pUAS-nucLacZ/+; Mef2-GAL4/+* |
| *Mef2TS > MECP2* | *yw; tub-Gal80^ts/+; Mef2-GAL4/pUAS-MECP2* |
| *how24B > MECP2* | *yw; +; how24B-GAL4/pUAS-MECP2* |
| *how24B > GFP* | *yw; pUAS-GFP/+; how24B-GAL4/+* |
| *vm > MECP2* | *yw; pUAS-MECP2/+; vm-GAL4/+* |
| *vm > GFP* | *yw; pUAS-GFP/+; vm-GAL4/+* |
| *c179 > MECP2* | *yw; pUAS-MECP2/+; c179-GAL4/+* |
| *c179 > GFP* | *yw; pUAS-GFP/+; c179-GAL4/+* |
| *hand > MECP2* | *yw; +; hand-GAL4/pUAS-MECP2* |
| *hand > GFP* | *yw; pUAS-GFP/+; hand-GAL4/+* |
| *hand > nLacZ* | *yw; pUAS-nucLacZ/+; hand-GAL4/+* |
| *tey > MECP2* | *yw; pUAS-MECP2/+; tey-GAL4/+* |
| *tey > GFP* | *yw; pUAS-GFP/+; tey-GAL4/+* |
| ***Neuronal drivers*** | |
| *Elav > MECP2* | *yw; Elav-GAL4; pUAS-MECP2* |
| *Elav > GFP* | *yw; Elav-GAL4; pUAS-GFP* |
| *gmr > MECP2* | *yw; gmr-GAL4; pUAS-MECP2* |
| *gmr > GFP* | *yw; gmr-GAL4; pUAS-GFP* |
| *D42 > MECP2* | *yw; D42-GAL4; pUAS-MECP2* |
| *D42 > GFP* | *yw; D42-GAL4; pUAS-GFP* |
| ***Enteric / gut drivers*** | |
| *Ilp > MECP2* | *w;;Ilp-GAL4; pUAS-MECP2* |
| *Ilp > GFP* | *w;;Ilp-GAL4; pUAS-GFP* |
| *Mip > MECP2* | *w;;Mip-GAL4/TM6B; pUAS-MECP2* |
| *Mip > GFP* | *w;;Mip-GAL4/TM6B; pUAS-GFP* |
| *NP3270 > MECP2* | *;NP3270-GAL4; pUAS-MECP2* |
| *NP3270 > GFP* | *;NP3270-GAL4; pUAS-GFP* |
| *NP3207 > MECP2* | *;NP3207-GAL4; pUAS-MECP2* |
| *NP3207 > GFP* | *;NP3207-GAL4; pUAS-GFP* |
| *mex > MECP2* | *w[1118]; P{w[+mC]=mex1-GAL4.2.1}10-8; pUAS-MECP2* |
| *mex > GFP* | *w[1118]; P{w[+mC]=mex1-GAL4.2.1}10-8; pUAS-GFP* |

**Table S1 List of genotypes used in the study.** The table reports, for each sample, the abbreviated genotype used throughout the study (left column) and the corresponding full genotype (right column).

| Driver | Tissue expression | survival |
|---|---|---|
| actin | whole body | 0% |
| elav | brain | 100% |
| D42 | nervous system | 96% |
| gmr | eye discs | 37% |
| ilp | anterior enteric neurons | 100% |
| Mef2 | body wall, visceral muscles, salivary glands | 0% |
| how24B | body wall, visceral muscles, trachea, salivary glands | 0% |
| c179 | body wall, visceral muscles, salivary glands | 17% |
| vm | body wall, visceral muscles | 0% |
| cg | fat bodies | 15% |
| Myo | enterocytes | 97% |
| NP3270 | gut (enterocytes) | 100% |
| NP3207 | gut (enterocytes) | 100% |
| mex | gut (enterocytes) | 100% |

**Table S2. Tissue expression patterns and adult survival of *Gal4* drivers used for tissue-specific *MECP2* misexpression.** Survival was calculated based on expected Mendelian ratios. Data represents the average of three independent biological replicates, each including approximately 50 animals.

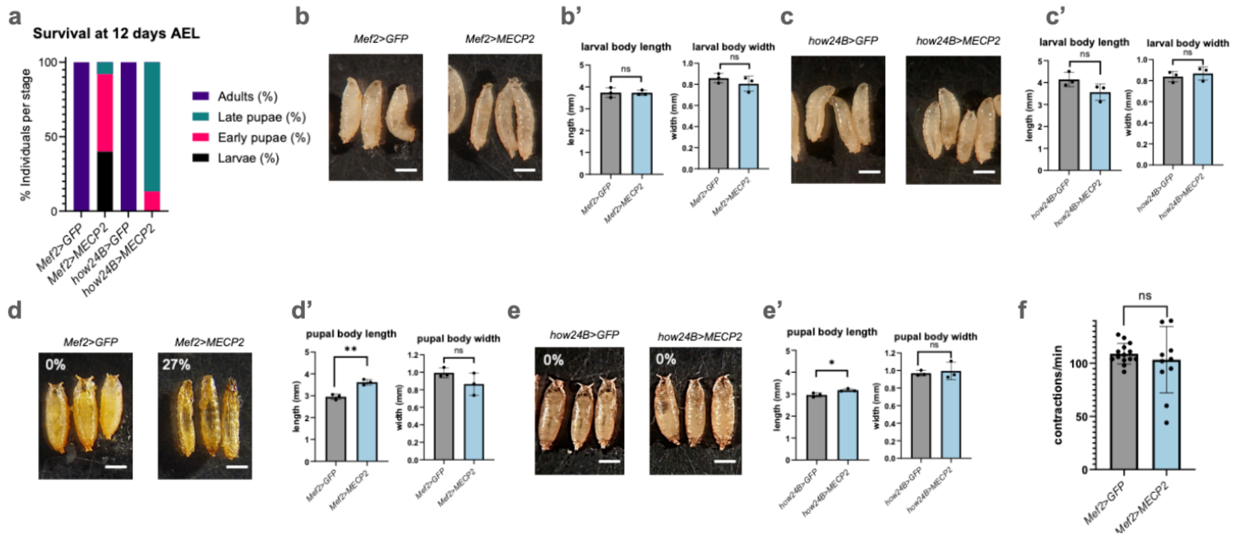


**Figure S1. Mesoderm-specific *MECP2* overexpression causes developmental delay and lethality. (a)** Stacked bar chart showing the % of animals at the indicated developmental stage at 12 days AEL for the indicated genotype. Each bar shows the mean of three independent biological replicates (N=50 per replicate). **(b-c)** Representative images of L3 larvae of the indicated genotypes. Scale bars: 1 mm. **(b'-c')** Quantification of larval length and width. Bar graphs represent the mean of three biological replicates (N = 30 animals/replicate). **(d-e)** Representative images of pupae for the indicated genotypes. Percentages indicate the occurrence of elongated larvae in figure, averaged across three biological replicates (N = 30 animals/replicate). Scale bars: 1 mm. **(d'-e')** Quantification of pupal length and width. Bar graphs represent the mean of three biological replicates (N = 30 animals/replicate). **(f)** Mouth hook contraction frequency of wandering L3 larvae from 4 independent biological replicates. In each replicate, 5 larvae were recorded and each data point represents the corresponding mean.

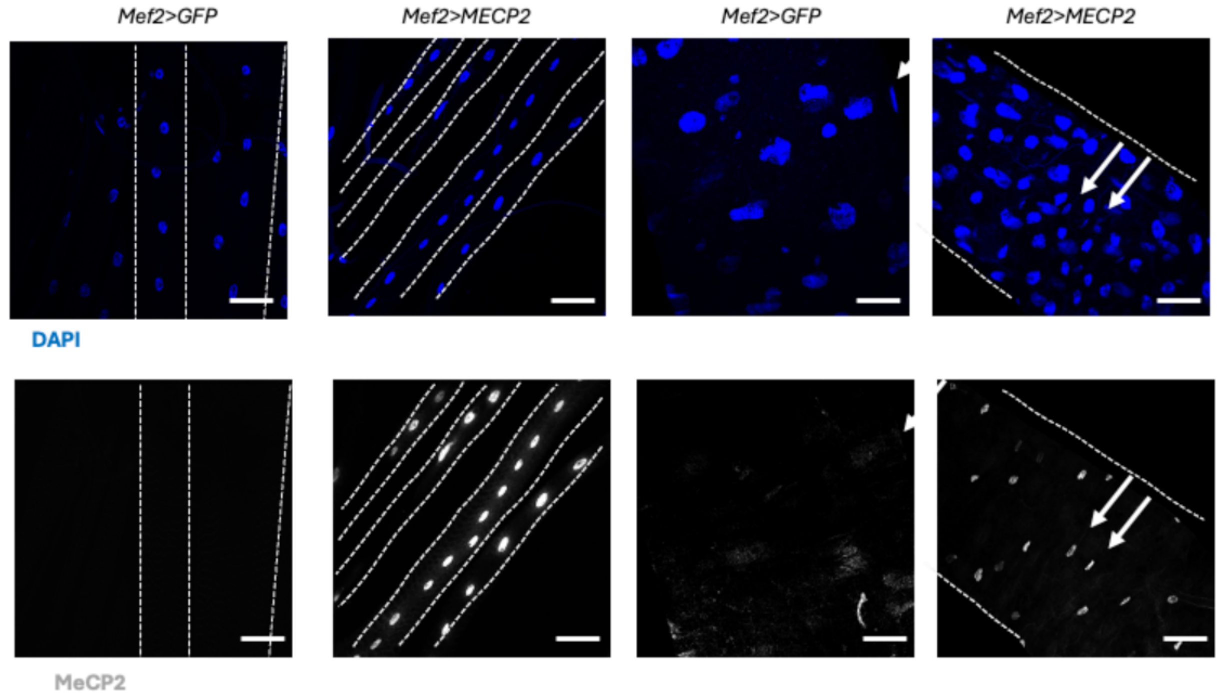


**Figure S2.** Immunostaining of nuclei with DAPI (blue) and MeCP2 (white) in larval body wall muscles (left) and gut (right). The arrows indicate the localization of MeCP2 in muscle cell nuclei. Scalebar: 50µm.

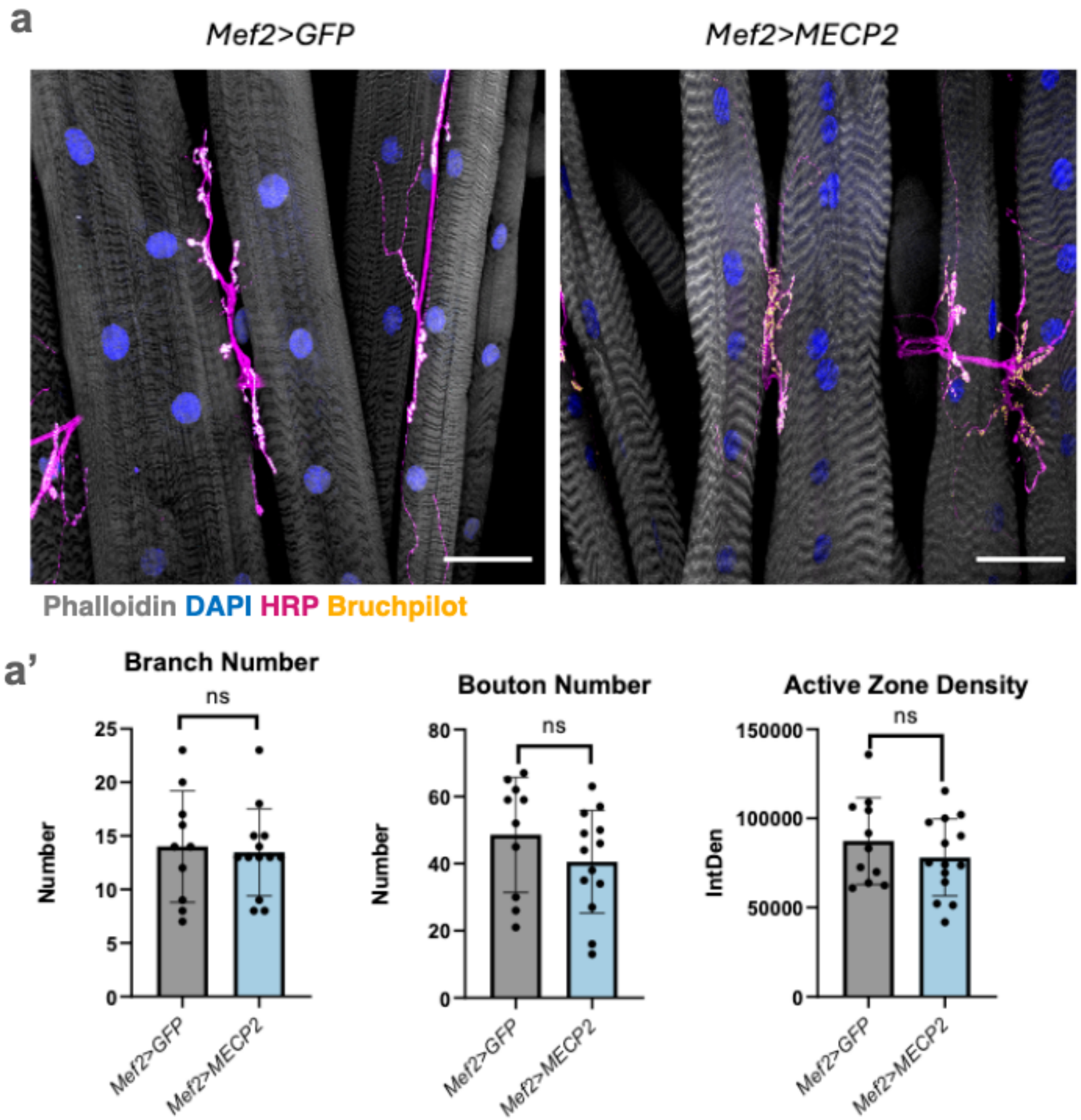


**Figure S3. Neuro-muscular junction morphology. (a)** Representative confocal images of larval NMJs and muscles stained with Phalloidin (grey), DAPI (blue), to detect HRP (magenta), a neuronal marker, and to detect Bruchpilot, a marker of active zones (yellow). Scale bar: 50 µm. **(a')** Quantification of NMJ branching, bouton number, and active zone density was performed as described in methods. Each point represents a single NMJ connecting muscles 6 or 7 in segments hemisegments A2–A4 (N ≥ 10 NMJs per genotype).

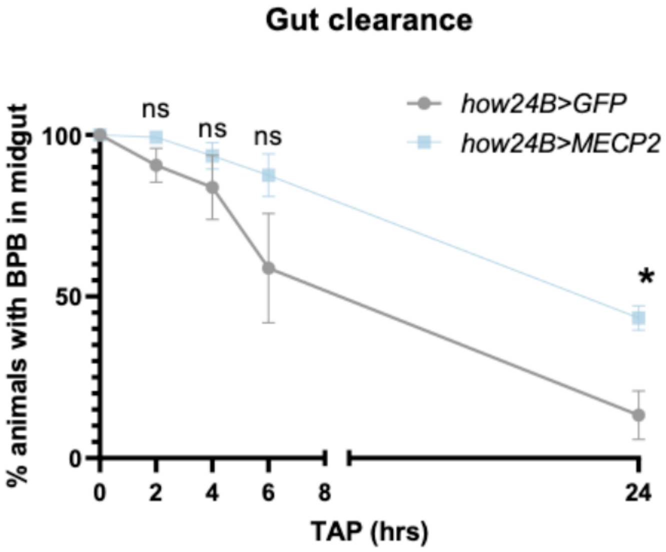


**Figure S4. Gut clearance assay with *how24B-GAL4* driver.** Graph representation of the % of larvae that retain dyed food over time. Control larvae efficiently cleared the food within 24 h prior to pupariation, indicating normal GI transit. In contrast, some *how24B>MECP2* larvae retained the colored food. Data are mean ± SEM. Approximately 30 larvae were analyzed per experiment, across 4 independent experiments.

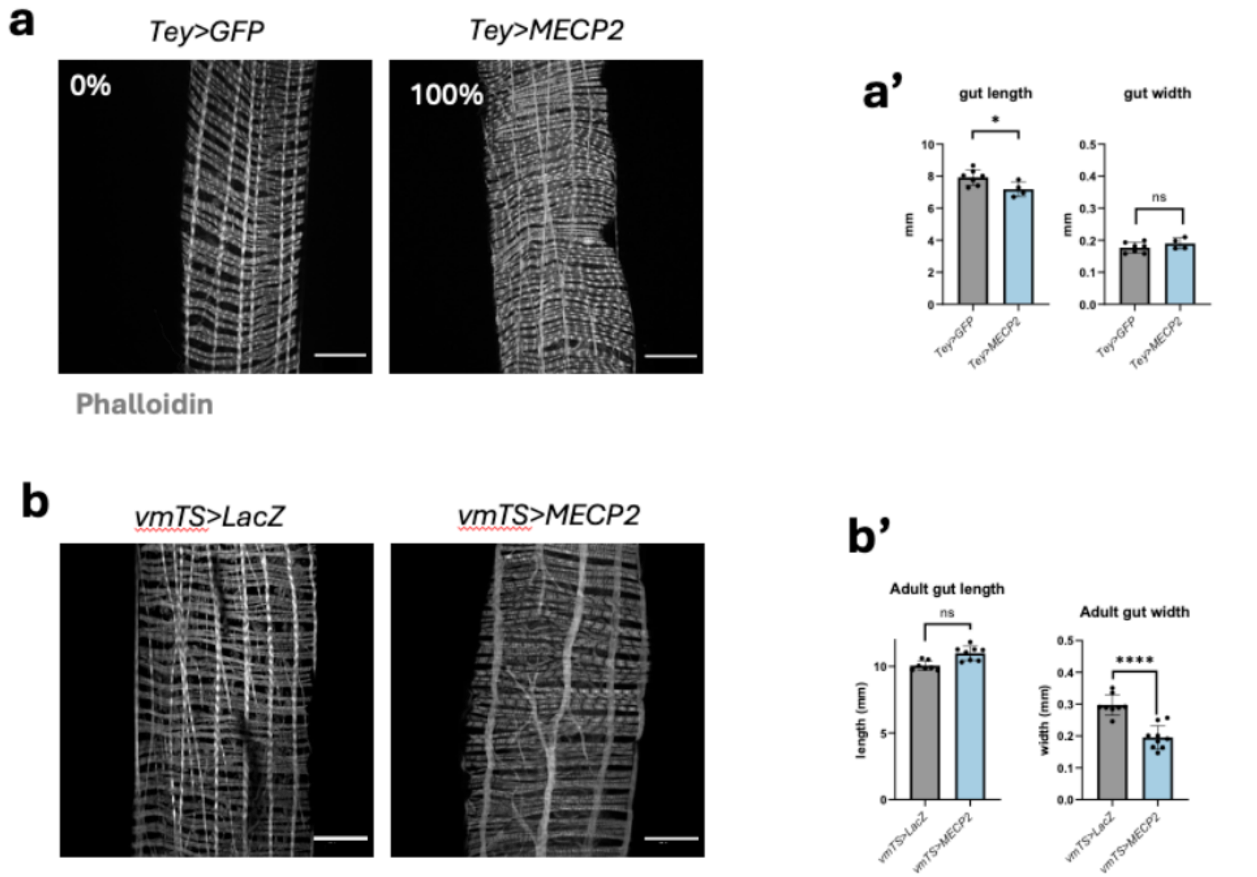


**Figure S5. Visceral muscle morphology. (a)** Representative confocal images of the visceral muscles of the indicated genotypes stained with phalloidin. *tey-Gal4* driver is specific for the longitudinal component of the visceral muscle. In *tey>MECP2* animals, the longitudinal fibers appear markedly thinned, a phenotype consistently observed in 100% of examined samples. Scale bar: 50 µm. (N=15 animals) **(a')** Quantification of gut length and gut width for the indicated genotypes. Each dot represents one gut. **(b)** Representative confocal images of *vmTS*-driven *MECP2* expression induced at 29 °C immediately after eclosion. A branching phenotype can be observed. Scale bar: 50 µm. **(c')** Quantification of gut length and width for the indicated genotypes. Each dot represents one gut (N=15 animals).

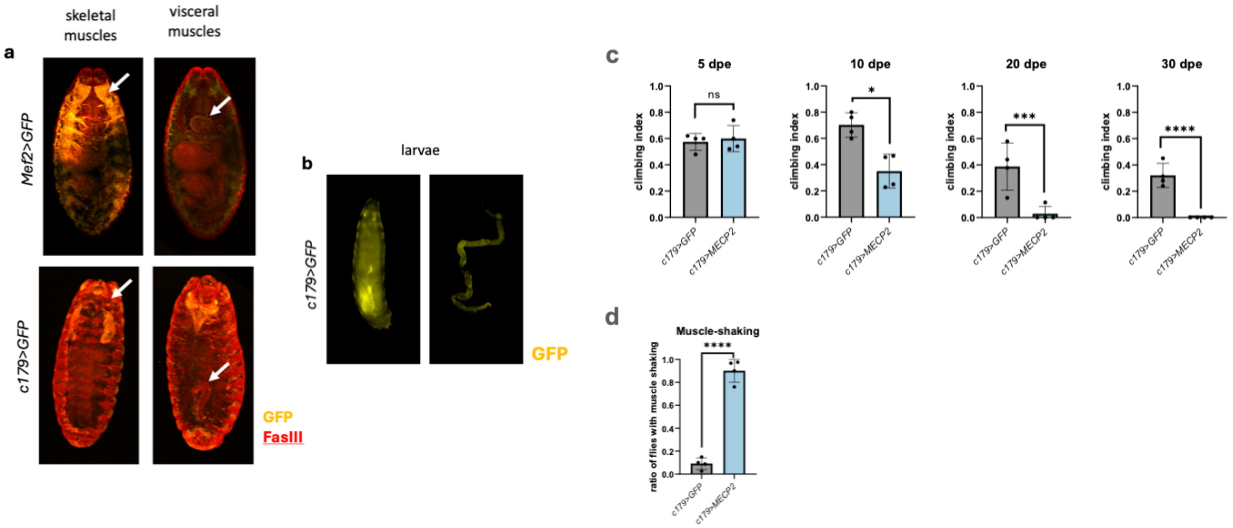


**Figure S6. Temporal expression of *c179>GFP* and *Mef2>GFP* in *Drosophila* embryos and larvae. (a)** Immunostaining of synchronized embryos with anti-GFP (yellow), anti-FasIII (red) to label cell membranes, and DAPI (blue). White arrows highlight skeletal muscles and visceral muscles; no GFP signal is detectable at this embryonic stage in *c179>GFP* embryo, only FasIII marks skeletal and visceral muscles (white arrows). N=50. Scalebar: 50µm **(b)** Representative *c179>GFP* L3 larvae, showing endogenous GFP fluorescence in skeletal and muscles (yellow), indicating that *c179* expression begins at the larval stage (N≥3). **(c)** Adult climbing assay on adults of the indicated genotypes. Performance was assessed at 5, 10, 20 and 30 dpe. Each data point represents the mean of 3 technical measurements, each performed on 10 flies, quantifying the fraction of animals that climbed 7 cm within 10 s. **(d)** Muscle shaking assessment in the indicated genotypes at 30 dpe. Each point represents the mean of 3 technical measurements, each performed on 10 flies, quantifying the fraction of animals displaying the shaking phenotype upon mechanical stimulation.

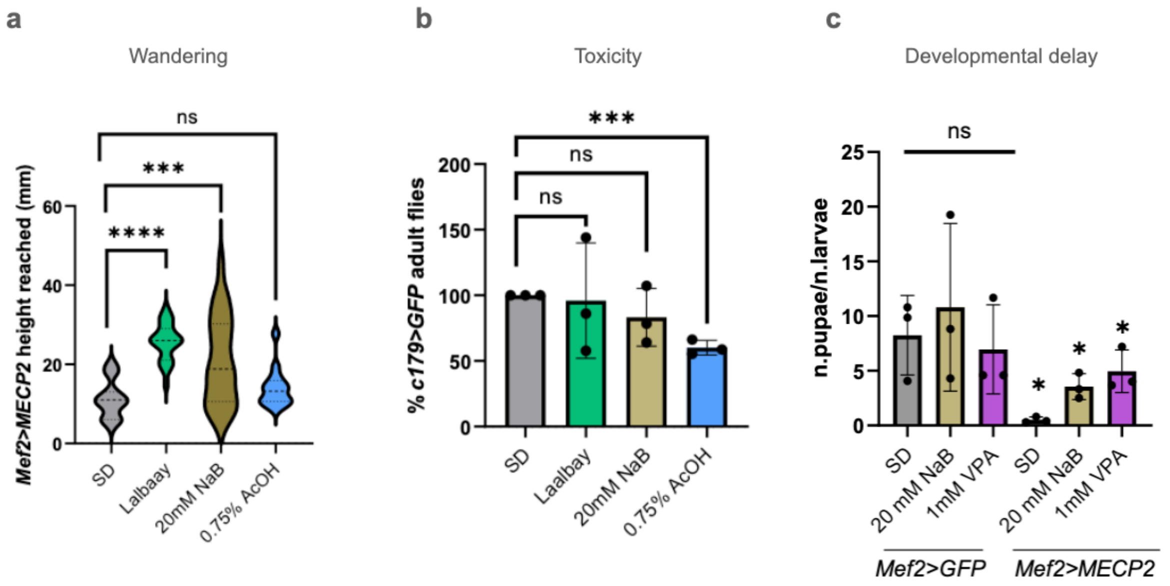


**Figure S7. (a)** Violin plot showing larval wandering ability, quantified as pupal height reached (in mm) as a readout of locomotor ability in larvae of the indicated genotype supplemented as indicated. Data from three independent crosses are pooled (N=20/replicate). **(b)** Bar graph showing adult survival of *c179>GFP* control animals, some supplemented as indicated. 3 independent biological replicates with N = 80 animals per replicate were perfomed. **(c)** Bar graph showing developmental delay of *Mef2>GFP* and *Mef2>MECP2* animals as described in methods. Each dot represents one biological replicate (N = 80 animals/replicate).

**Supplementary videos can be accessed at:**

https://drive.google.com/drive/folders/13XrtKZ47vbSP4rCMVFPYQ6SPx7tKGKNI?usp=sharing

**Video S1. Mouth hook contraction assay**. **(a-b)** Representative recordings of a technical replicate consisting of five third-instar larvae from the same cross, placed in PBS supplemented with yeast. The video shows the rhythmic mouth hook movements used to quantify larval feeding activity. Mouth hook contractions were counted for each larva over a 1-minute interval, as described in methods.

**Video S2. Larval gut contraction assay. (a-d)** Representative recordings showing gut peristaltic contractions in Schneider's *Drosophila* medium for each genotype and treatment. The anterior acidic region of the gut is highlighted in yellow, and the remaining gut in blue, reflecting feeding with blue dye in the food. The right side of the video shows the aboral region of the gut. Gut movements were quantified as described in methods. Each recording corresponds to a technical replicate.

**Video S3. Representative video showing the muscle shaking phenotype in aged flies.** Thirty-day-old adult flies of the indicated genotypes (*c179>GFP, c179>MECP2*) were placed in transparent vials as labelled. The video shows a representative example of the muscle shaking phenotype observed upon stimulation.
